# Invariance of object detection in untrained deep neural networks

**DOI:** 10.1101/2022.09.08.507096

**Authors:** Jeonghwan Cheon, Seungdae Baek, Se-Bum Paik

**Author notes:** **Correspondence:** Se-Bum Paik.

## Abstract

The ability to perceive visual objects with various types of transformations, such as rotation, translation, and scaling, is crucial for consistent object recognition. In machine learning, invariant object detection for a network is often implemented by augmentation with a massive number of training images, but the mechanism of invariant object detection in biological brains — how invariance arises initially and whether it requires visual experience — remains elusive. Here, using a model neural network of the hierarchical visual pathway of the brain, we show that invariance of object detection can emerge spontaneously in the complete absence of learning. First, we found that units selective to a particular object class arise in randomly initialized networks even before visual training. Intriguingly, these units show robust tuning to images of each object class under a wide range of image transformation types, such as viewpoint rotation. We confirmed that this “innate” invariance of object selectivity enables untrained networks to perform an object-detection task robustly, even with images that have been significantly modulated. Our computational model predicts that invariant object tuning originates from combinations of non-invariant units via random feedforward projections, and we confirmed that the predicted profile of feedforward projections is observed in untrained networks. Our results suggest that invariance of object detection is an innate characteristic that can emerge spontaneously in random feedforward networks.

**Highlights:**

- Object-selective units spontaneously emerge in untrained deep neural networks
- Object selectivity maintains robustly in a wide range of image transformations
- Feedforward model can explain spontaneous emergence of the invariance
- Innate invariance enables invariant object detection without learning to variations

## 1 Introduction

Visual object recognition is a crucial function for animal survival. Human and primates can detect objects robustly, despite huge variations in the position, size, and viewing angles (Logothetis et al., 1994; Tanaka, 1996; Connor et al., 2007; Pinto et al., 2008; DiCarlo et al., 2012; Poggio and Ullman, 2013). This challenging ability is thought to be based on invariant neural tuning in the brain — Neurons that selectively respond to various objects have been observed in higher visual areas, and these neurons showed invariant object representation across various types of transformation (Perrett et al., 1991; Ito et al., 1995; Wallis and Rolls, 1997; Hung et al., 2005; Zoccolan et al., 2007; Li et al., 2009a; Freiwald and Tsao, 2010; Apurva Ratan Murty and Arun, 2015; Ratan Murty and Arun, 2017). Behavioral- and neural-level observations of this function have led many researchers to raise the important question of how this invariance of object detection emerges.

Often, this invariant neural tuning has been considered to develop from the learning of various types of visual transformations (Biederman, 1987; Li et al., 2009b). With the notion that visual experience of natural objects contains numerous variants that transform depending on the viewing conditions, it has been suggested that the capability to detect objects invariantly can develop gradually when observers repeatedly see objects with a wide range of variations (Földiák, 1991). Notably, in the machine learning field, invariant object recognition is also implemented via learning with a massive dataset. In this case, data augmentation, a specialized method to increase the dataset volume, is often applied (Simard et al., 2003; O’Gara and McGuinness, 2019; Shorten and Khoshgoftaar, 2019) to generate images through linear transformations such as rotation, positional shifting, and flipping, as in natural visual experience. Then, invariant object recognition is achieved from the training of the augmented dataset with computer vision models (Chen et al., 2019; O’Gara and McGuinness, 2019; Shorten and Khoshgoftaar, 2019). In contrast to the above scenario, observations in newborn animals suggest the possibility of its emergence without learning: Human infants show a preference to faces despite variations of the size and rotations in depth (Turati et al., 2008; Kobayashi et al., 2012; Ichikawa et al., 2019). In addition, newborn chicks can detect virtual objects from novel viewpoints (Wood, 2013). These findings imply that invariant object detection arises without visual experience, but the developmental mechanism of this invariance in biological brains – how object invariance arises innately in the complete absence of learning – remains elusive.

A model study using a biologically inspired deep neural network (DNN) (Krizhevsky et al., 2012; Simonyan and Zisserman, 2015) has been suggested as an effective approach to this problem (Paik and Ringach, 2011; DiCarlo et al., 2012; Yamins and DiCarlo, 2016; Sailamul et al., 2017; Baek et al., 2020, 2021; Jang et al., 2020; Kim et al., 2020, 2021; Park et al., 2021; Song et al., 2021). DNNs consist of a stack of feedforward projections inspired by the hierarchical structure of the visual pathway and can be used as simplified model to investigate various visual functions. For instance, it was reported that a DNN trained to natural images can predict neural responses in the primate visual pathways from an early visual area (e.g., primary visual cortex, V1) to a higher visual area (e.g. inferior temporal cortex, IT) (Cadieu et al., 2014; Yamins et al., 2014). Recent studies also provided insight into the origin of functional tuning in the brain, by showing that units that selectively respond to numerosity, faces, and various types of objects among visual stimuli can arise in a randomly initialized DNN without any learning (Baek et al., 2021; Kim et al., 2021).

By adopting a similar approach, here we show that object invariance can arise in completely untrained neural networks. Using AlexNet (Krizhevsky et al., 2012), a model designed along the structure of the visual stream, we found that units selective to various visual objects are observed in a randomly initialized DNN and that these units maintain selectivity across a wide range of variations, such as the viewpoint, even without any visual training. We observed that a certain proportion of the units show an invariantly tuned viewpoint, while other groups of units show tuning to a specific viewpoint. Preferred feature images obtained from the reverse-correlation method showed that each specific viewpoint unit encodes a shape from a particular view of an object, while invariant viewpoint units encode inclusive features from specific units with different preferred angles. We found that invariant units emerge by homogenous projections from specific units in the previous layer in a random feedforward network. Finally, we confirmed that this innate invariance enables the network to perform an object-detection task under an enormous range of variations of viewpoints. Overall, our results suggest that invariant object detection can emerge spontaneously from the random wiring of hierarchal feedforward projections in an untrained deep neural network.

## 2 Results

### 2.1 Emergence of object selectivity in untrained networks

To investigate the emergence of invariant object selectivity in an untrained model network, we used AlexNet (Krizhevsky et al., 2012), a biologically inspired DNN that models the structure of the ventral visual pathway. To find an object-selective response of an individual unit in the network, we investigated the responses of the final convolutional layer (Conv5), which is presumed to correspond to the IT domain of the brain. To simulate the condition of an untrained hierarchical network, we randomly initialized AlexNet using a standardized network initialization method (LeCun et al., 1998), by which the weights of the filters in every convolutional layer are randomly selected from a Gaussian distribution.

The stimulus set was designed to contain nine different object categories (e.g. Monitor, Bed, Chair, etc.) (**Figure 1A**). To define selective units for a specific target object, eight other class object sets and one scrambled set of the target object were used, following a previous experimental study (Stigliani et al., 2015) (see Methods for details). The images in each class were prepared by controlling the low-level features of the luminance, contrast, object location and object size (**Supplementary Figure 1**). Specifically, the pixel value distribution of the object image and background image were calibrated using the same Gaussian distribution (mean = 127.5, s.d. = 51.0), and the intra-class image similarity was also controlled at statistically comparable level. As this stimulus was given as input for the randomly initialized networks, the responses were measured in the Conv5 layer and an analysis of object selectivity was conducted (**Figure 1B**).

**Figure 1.**
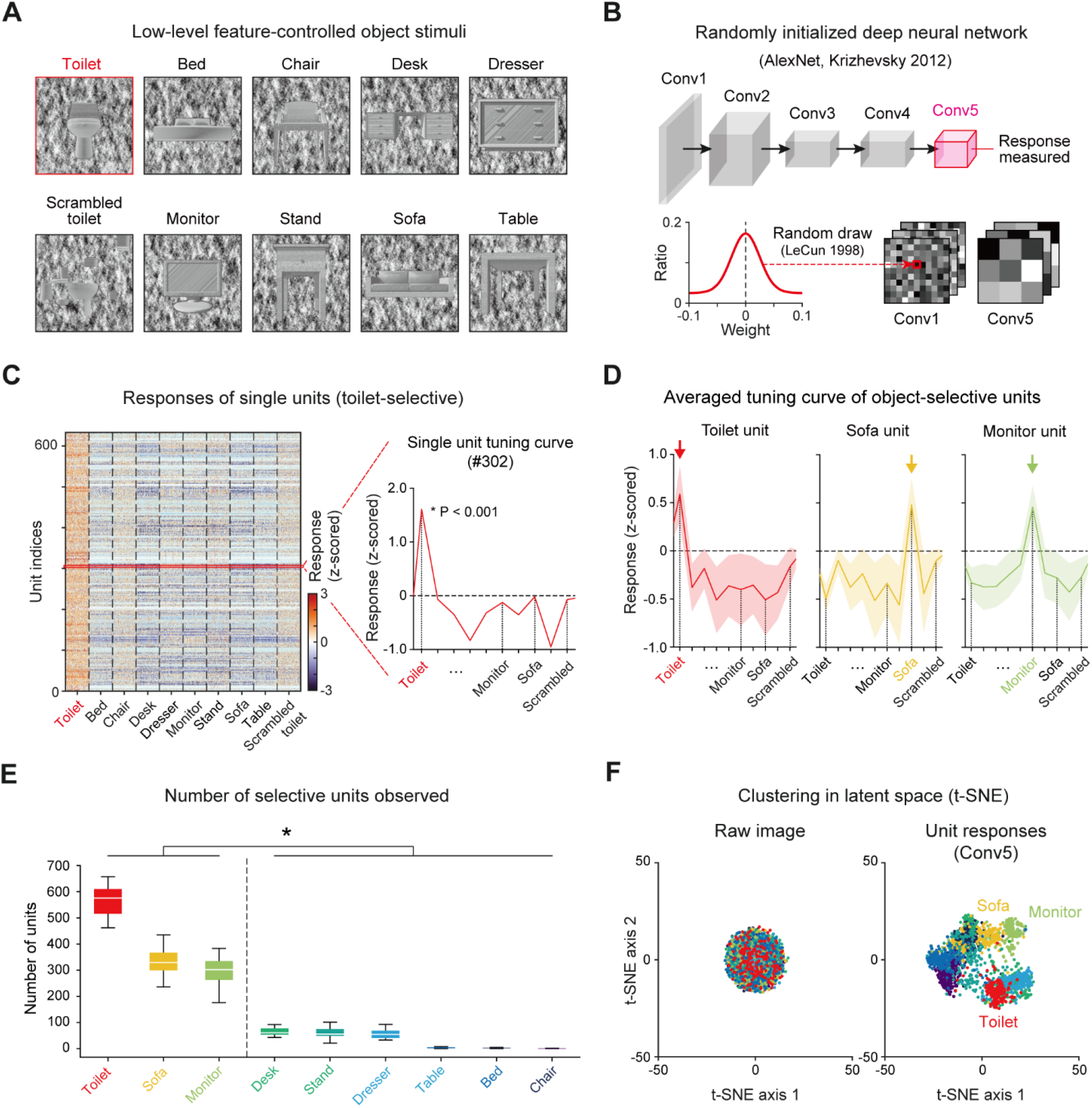
Emergence of selectivity to various objects in untrained networks: **(A)** The stimulus images were selected and modified from a publicly available CAD dataset [https://modelnet.cs.princeton.edu/] (See Methods for details). The images contain nine different object categories. The low-level features of the luminance, contrast, object location and object size of images were calibrated equally across object classes. **(B)** The structure of a randomly initialized deep neural network. Five convolutional layers in AlexNet (Krizhevsky et al., 2012) were randomly drawn from a Gaussian distribution (LeCun et al., 1998). **(C)** Responses of each single toilet-selective unit (P < 0.001, two-sided rank-sum test). The red curve is an example tuning curve of a single unit. **(D)** Average responses of selective units for three object classes (Toilet, n = 548; Sofa, n = 285; Monitor, n = 301). Each arrow indicates the preferred object class. Shaded areas represent the standard deviation from the tuning curves of the target units. **(E)** The number of object-selective units for nine classes in untrained networks (n = 20). Box plots indicate the inter-quartile range (IQR between Q1 and Q3) of the dataset, the white line depicts the median and the whiskers correspond to the rest of the distribution (Q1 − 1.5*IQR, Q3 + 1.5*IQR). **(F)** Visualization of the latent space by the t-SNE method (Maaten and Hinton, 2008) from raw images and the responses of Conv5 units to each class. The raw images of each object class do not cluster in the latent space, but the responses of the untrained network collected in Conv5 were clustered in the latent space according to the class of the given image.

We found object-selective units that show higher responses to a specific class of target images (e.g. Toilet) than to other non-target class images and scrambled images (P < 0.001, two-sided rank-sum test) (**Figure 1C**). Among the nine object categories, we observed that object-selective units emerge mostly in a few objects categories (**Figures 1D and E**). We observed toilet-selective units (n = 565 ± 55 in 20 random networks, mean ± s.d.), sofa-selective units (n = 339 ± 68), and monitor-selective units (n = 294 ± 54) in the Conv5 layer (43,264 units; 13 × 13 × 256, *N*_x-position_ × *N*_y-position_ × *N*_channel_) (**Figure 1D**). In particular, the number of object-selective units was divided into large and small groups (**Figure 1E**, n = 20, two-sided rank-sum test, *P < 10^−27^). Large groups consisted of the toilet, sofa, and monitor groups (n_units_ = 400 ± 133) and small groups were the dresser, desk, bed, chair, nightstand, and table groups (n_units_ = 32 ± 32). In the subsequent analyses, we investigated the results mostly for the three object classes which show a large number of selective units.

We investigated the number and the selective indexes of object units across the convolutional layers and found that the number of object units increases when the convolutional layers become deeper (**Supplementary Figure 2A**). The object-selective index for a single unit also shows a strong tendency to increase across convolution layers, demonstrating that object tuning becomes sharper through the network hierarchy (**Supplementary Figure 2B**). Furthermore, we found that the responses of an untrained network measured in the deep layer (Conv5) were clustered as object classes in the latent space, while raw images do not cluster in the latent space (**Figure 1F**).

### 2.2 Invariance of object-selective units in untrained networks

Next, to investigate whether the observed object-selective units show viewpoint-invariant representations of an object image, we measured the responses of object-selective units to target objects and non-target objects with various viewpoint angles. To do this, a viewpoint-variant stimulus set was generated (**Supplementary Figure 3**) by rotating the viewpoint of 3D objects on the horizontal plane (**Figure 2A**). For each object, we rendered 13 variant images at different viewpoints between −90° and 90°. Then, we measured the responses of selective units to target objects and non-target objects with various viewpoint angles (**Figure 2B**). We found that units show selective responses when an object image within a certain threshold is presented (**Figure 2C, left**, Toilet at 0° vs. Non-toilet, *P < 10^−13^; Toilet at 45° vs. Non-toilet, **P < 10^−5^), while the units did not show selectivity when an object image at a larger viewpoint angle was given (**Figure 2C, left**, Toilet at 90° vs. Non-toilet, n = 200, one-sided rank-sum test, NS, P = 0.492). Hence, the selectivity of object-selective units is maintained within a limited effective range (**Figure 2C, right**).

**Figure 2.**
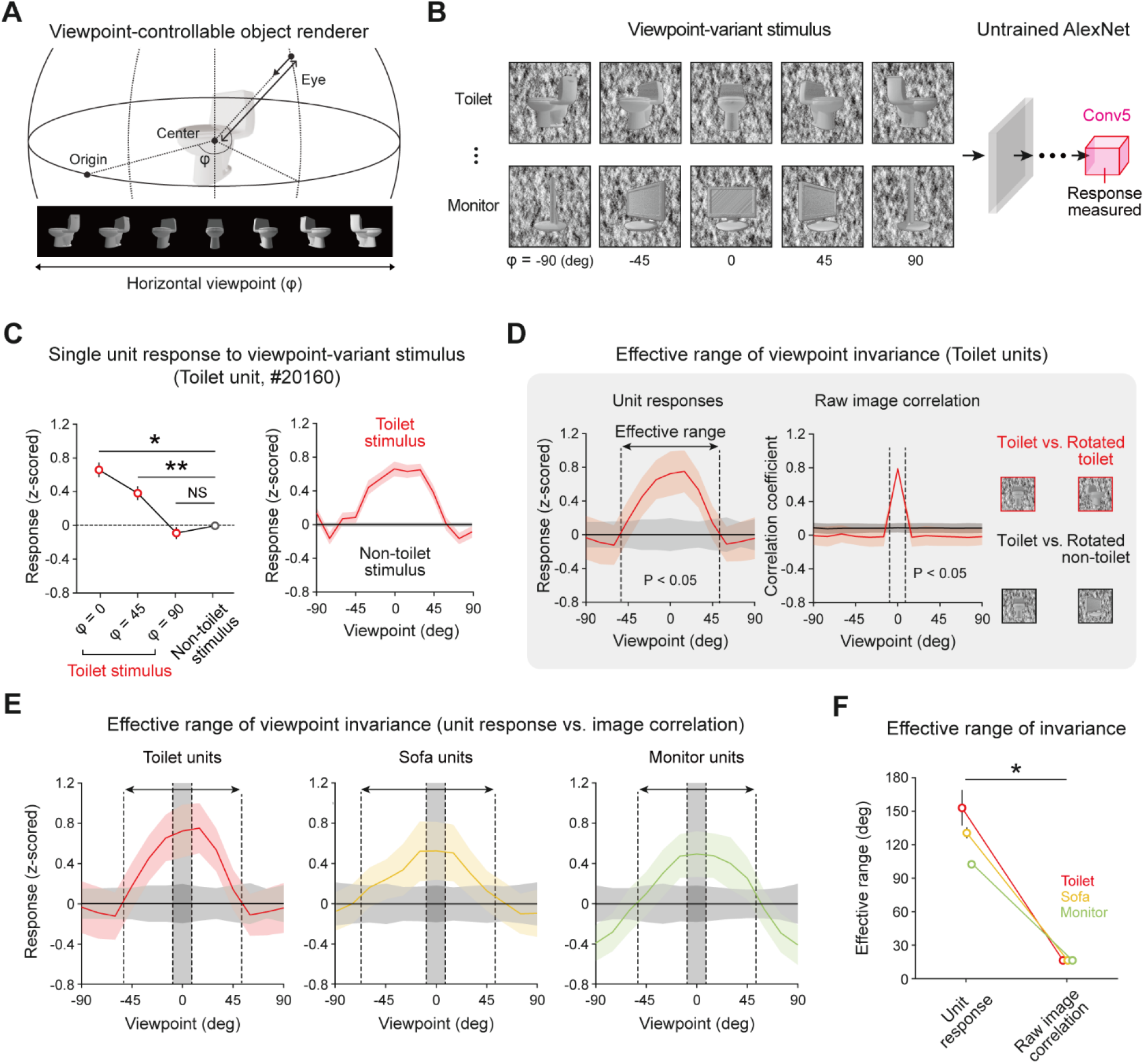
Viewpoint-invariant object selectivity observed in an untrained network: **(A)** An object renderer generates object images at various viewpoints rotating in a horizontal orbit. **(B)** The viewpoint-varying object stimulus was generated within the viewpoint range of −90° to +90° with 13 steps. The responses of the object-selective unit were measured in the final convolutional layer of an untrained AlexNet. **(C)** Viewpoint-invariant responses of a selective unit for viewpoint-rotated object stimulus; the tuning of a single toilet-selective unit shows a wide range of viewpoint invariance. Shaded areas and error bars represent the standard error of 200 images. **(D)** The effective range of the average responses of object units and the pixel-wise correlation of a raw image (n = 200, one-sided rank-sum test, P < 0.05). **(E)** Average response of object-selective units for an object stimulus at different viewpoint rotations. The arrow indicates the effective range of the selective response, and the shaded area indicates the effective range of the raw-image correlation. Shaded areas and error bars represent the standard deviation of 200 images. **(F)** Comparison of effective ranges between the selective response and raw-image correlation in each object-selective unit. Error bars indicate the standard deviation of 20 random networks.

To investigate the effective range that maintains the selectivity of object units quantitatively, we investigated the responses of selective units with a viewpoint between −90° and 90° and estimated the boundary of the viewpoint variation around which target-object tuning is lost. For example, we observed that object tuning of toilet units was retained when the viewpoint change was within 105° (**Figure 2D, left**, n = 200, one-sided rank-sum test, P < 0.05). Then, to verify whether the viewpoint invariance of an object-selective unit simply arises due to the similarity of the object shape upon a change of the viewpoint, we estimated the pixel-wise raw-image correlations between object images from a front view and a rotated view. We compared the effective ranges of viewpoint invariance between the selective responses and the image correlations (**Figure 2D, right**). For toilet units, we observed that the effective range of the selective responses is significantly wider than that of the image correlation (**Figure 2E**, Toilet units). Similarly, this tendency was commonly observed in other object-selective units (**Figures 2E and 2F**, n = 20, two-sided rank-sum test; toilet unit, *P < 10^−4^; sofa unit, *P < 10^−4^; monitor unit, *P < 10^−4^). This result suggests that the observed invariance is not simply due to the similarity of the object images at different viewpoints but is a characteristic of object-selective units in untrained networks. To find the origin of the invariance in an untrained network, we also examined the single-unit-level characteristics of invariance. We found that each unit shows considerable variations in the response characteristics when a target object image with various viewpoints is given as the input. In particular, each unit shows various effective ranges (**Figures 3A and 3B, left**). Considering the definition of viewpoint invariance, we presumed that the top 30% of units were “viewpoint-invariant” units and the bottom 30% units were “viewpoint-specific” units in the subsequent analyses. Indeed, we observed that each tentative viewpoint-specific unit has various preferred angles; i.e., they only respond to a particular view of an object (**Figure 3B, right**).

**Figure 3.**
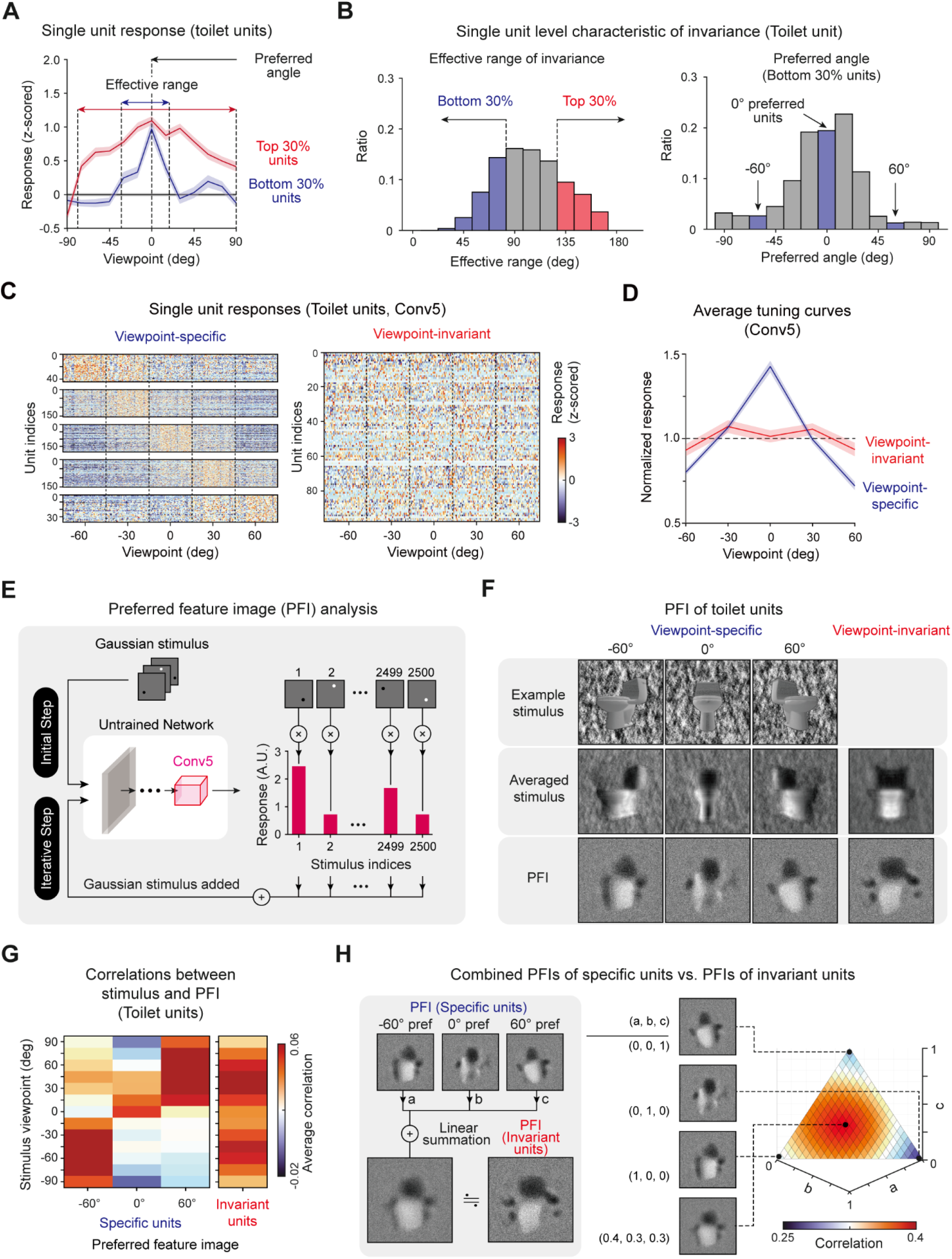
Single-unit-level analysis of invariance: **(A)** Viewpoint tuning curves of two sample toilet units: a unit with a wide effective range (Top 30%) and a unit with a narrow effective range (Bottom 30%). **(B)** Histogram of the invariance effective range of each unit and histogram of the preferred angle of units with a narrow effective range (Bottom 30%). **(C)** Responses of individual viewpoint-specific and viewpoint-invariant toilet-selective units in the Conv5 layer (viewpoint-specific units, one-way ANOVA, P < 0.05; viewpoint-invariant units, one-way ANOVA, P > 0.05). **(D)** Average tuning curves of viewpoint-specific units (n = 147) and viewpoint-invariant units (n = 95) in an untrained network. Shaded areas represent the standard error of each type of unit. **(E)** Overall process of the preferred feature image (PFI) of target units in Conv5 of untrained networks using a reverse-correlation analysis (Bonin et al., 2011; Baek et al., 2021). The input stimulus was generated with a randomly positioned bright or dark dot blurred with a 2D Gaussian filter. The PFI was calculated as the response-weighted summation of the input stimulus. **(F)** Obtained preferred feature images of viewpoint-invariant and viewpoint-specific units with different preferred angles (−60°, 0°, 60°). An example stimulus and an average stimulus corresponding to preferred viewpoint angle. For invariant units, the average stimulus across all viewpoints is presented. **(G)** Correlation between the stimulus of various viewpoints and PFIs of various types of units. **(H)** Comparison between the PFI of an invariant unit and the weighted summation of PFIs of specific units. Here, ‘a’, ‘b’ and ‘c’ represent the weight of each PFI for summation (a + b + c = 1). The 3D plot represents the pixel-wise correlations for different values of each weight pairs.

To verify our conjecture that “viewpoint-specific” and “viewpoint-invariant” units exist and can be classified according to the observed effective range of each unit (**Figures 3A and 3B**), we investigated the responses of object-selective units for object images with different viewpoints. Target object images with a viewpoint between −60° and −60° (five steps, 50 images per viewpoint class) were presented to the network, and the responses were measured. Indeed, we observed that there are units that only respond to a particular viewpoint image (**Figure 3C, viewpoint-specific**, one-way ANOVA, P < 0.05) and units that respond invariantly to any viewpoint image (**Figure 3C, viewpoint-invariant**, one-way ANOVA, P > 0.05). Viewpoint-specific units show highly tuned responses to one preferred viewpoint angle, while viewpoint-invariant units show a flat tuning curve to any viewpoint (**Figure 3D**).

Next, to visualize the distinct tuning features of viewpoint-specific and viewpoint-invariant units, we used a reverse-correlation method (Bonin et al., 2011; Baek et al., 2021) and obtained the preferred feature images (PFIs) of units (**Figure 3E**, see Methods for details). We found that each specific unit showed a PFI similar to an object image at the viewpoint angle of its preferred value. From this result, we confirmed that each specific unit encodes a shape from a particular view of the object (**Figure 3F, Specific**). In contrast, the PFIs of viewpoint-invariant units were similar to the average stimulus image of various viewpoints (**Figure 3F, Invariant**). Notably, the calculation of the correlations between the stimulus of various viewpoints and the PFIs from each different type of unit reveals that the PFI of specific units shows a high correlation only with the stimulus image of the corresponding viewpoint, while that of invariant units shows high correlations with the stimulus images of various viewpoints (**Figure 3G**). From this observation, we hypothesized that the PFIs of invariant units can be expressed as a linear combination of the PFIs of specific units. We tested this scenario by searching for wiring coefficients that maximize the correlation between the PFIs of invariant units and a combined stimulus image (**Figure 3H, left**). We observed a very high correlation when each PFI of a specific unit is linearly combined with fairly homogeneous coefficients (**Figure 3H, right**). The same tendency was observed in the PFIs of other object-selective units (**Supplementary Figure 4**). These results suggest that viewpoint-invariant units can originate from a homogenous combination of viewpoint-specific units.

### 2.3 The feedforward model can explain the spontaneous emergence of invariance

To validate the hypothesis that viewpoint-invariant units originate from the projection of viewpoint-specific units in the previous layer, we backtracked projections of the units from the source layer (Conv4) to the target layer (Conv5) and examined the weights of connected viewpoint-specific units. First, we confirmed that viewpoint-specific (n = 765 ± 102) and viewpoint-invariant toilet-selective units (n = 130 ± 28) exist in Conv4 as well as in Conv5 (viewpoint-specific, n = 504 ± 81, viewpoint-invariant, n = 96 ± 16). We confirmed that the viewpoint-specific units in Conv5 receive stronger input from units with the same object tuning than from other units in Conv4 (**Figure 4A, left and middle**, n = 20, two-sided rank-sum test, *P < 10^−7^). In more detail, the viewpoint-specific units in Conv5 receive inputs from Conv4 units strongly biased to a particular viewpoint angle (**Figure 4A, right**, n = 20, one-way ANOVA, *P < 10^−11^). This tendency of a strongly biased weight also appeared in other preferred viewpoints. The connectivity between viewpoint-specific units with the same preferred angle in the source and target layers showed significantly high weights compared to other projection directions (**Figure 4B**, n = 20, one-way ANOVA, *P < 0.05).

**Figure 4.**
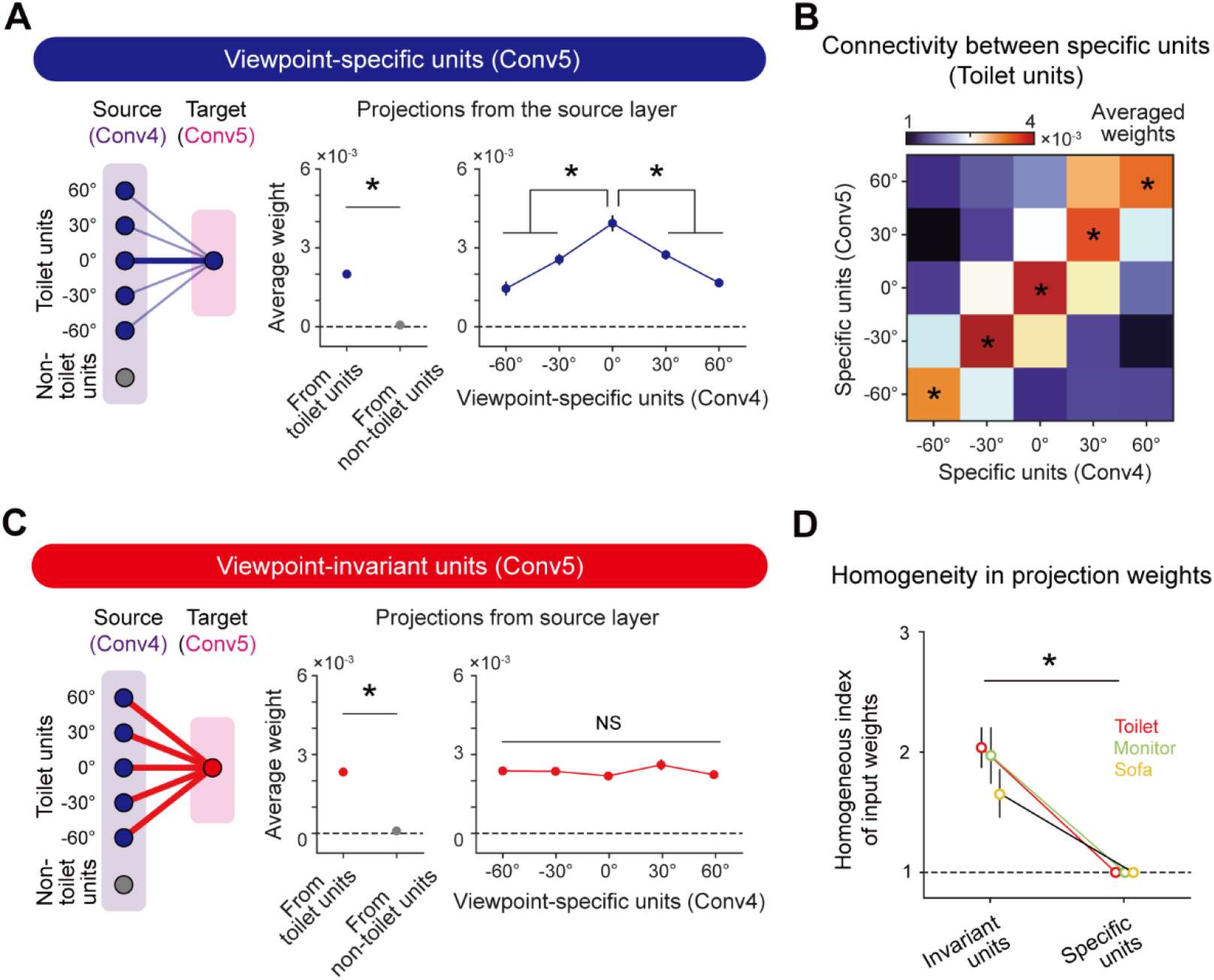
Emergence of viewpoint invariance based on unbiased projection from viewpoint-specific units: **(A)** Connectivity diagram (left); averaged weight from target object units and non-target object units (Conv4) to viewpoint-specific units (Conv5) (middle); averaged weight from viewpoint-specific units (Conv4) to viewpoint-specific units (Conv5). Error bars indicate the standard error of 20 random networks. **(B)** Heatmap of weights between specific units from the source layer and specific units from the target layer. This heatmap shows biased input to specific units. **(C)** Connectivity diagram (left); averaged weight from target object units and non-target object units (Conv4) to viewpoint-invariant units (Conv5) (middle); and averaged weight from viewpoint-specific units (Conv4) to viewpoint-invariant units (Conv5). Error bars indicate the standard error of 20 random networks. **(D)** The homogeneous index of the input projection weight that connected viewpoint-specific units (Conv4) to viewpoint-invariant units (Conv5). Error bars indicate the standard deviation of 20 random networks.

We also found that viewpoint-invariant units in Conv5 are strongly connected to units with the same object selectivity in Conv4 (**Figure 4C, left and middle**, n = 20, two-sided rank-sum test, *P < 10^−7^), as in the case of viewpoint-specific units. However, the viewpoint-invariant units in Conv5 receive homogeneous inputs from specific units in Conv4 units with various preferred viewpoint angles (**Figure 4C, right**, n = 20, one-way ANOVA, NS, P = 0.313). To estimate the degree of homogeneity in the projection weights, we defined the homogenous index as the inverse of the standard deviation of the average weight connected to specific units with different viewpoints in the source layer. The index of the average weight connected to viewpoint-invariant units is significantly higher than that of the average weight connected to the viewpoint-specific units, indicating an unbiased input to the viewpoint-invariant units (**Figure 4D**, toilet; invariant units vs. specific units, n = 20, two-sided rank-sum test, *P < 0.05). This tendency was also observed in units with other object tunings (**Figure 4D**, sofa and monitor; invariant units vs. specific units, n = 20, two-sided rank-sum test, *P < 0.05; **Supplementary Figure 5)**. This implies that observed viewpoint invariance of object tuning can originate from hierarchical random feedforward projections.

To verify this developmental model further, we revisited earlier observations of invariant object tuning in the monkey IT which reported that neurons in the higher layer in the hierarchy show increased invariance (from the middle lateral (ML) to the anterior medial (AM) area) (Freiwald and Tsao, 2010). We investigated whether this trend of invariance across layers is also observed in our model neural network, finding that such layer-specific characteristics of viewpoint invariance also emerge in the untrained network we used.

We observed that the level of invariance increased along the network hierarchy (**Supplementary Figure 6A**). To quantify these invariance characteristics, we introduced an invariance index of units, defined as the inverse of the standard deviation of responses across different viewpoints. We observed an increase in the invariance index of selective units higher up in the hierarchy in the untrained AlexNet (**Supplementary Figures 6B**). The viewpoint-invariance index in Conv4 is significantly higher than that in Conv3 (n = 20, two-sided rank-sum test, *P < 10^−7^). Also, the viewpoint-invariance index in Conv5 is significantly higher than that in Conv4 (n = 20, two-sided rank-sum test, **P < 10^−7^). This increasing tendency of the viewpoint-invariance index along the network hierarchy is also observed in other object-selective units (n = 20, two-sided rank-sum test; sofa, *P < 10^−7^, **P < 10^−7^; monitor, *P < 10^−7^, **P < 10^−7^). In addition, we confirmed that the same increasing tendency of the number of invariant units along the network hierarchy exists across various object tunings (**Supplementary Figure 6C**, n = 20, two-sided rank-sum test; toilet, *P < 0.05, **P < 0.001; sofa, *P < 0.05; monitor, *P < 10^−5^, **P < 10^−6^). These results suggest that our model provides a plausible scenario for understanding the spontaneous emergence of invariant object selectivity in untrained networks, which is supported by previous experimental observations of neural tunings.

### 2.4 Innate invariance enables invariant object detection without data-augmented learning

Next, we tested whether this innate invariance in untrained networks enables the network to perform the invariant object-detection task without learning. We expected that the information given by invariant object units is sufficient to detect an object while the viewpoint of the given object image varies, and in particular, that viewpoint-invariant units play a key role in enabling invariant object detection. To confirm this hypothesis, we designed two different methods to train an SVM which classifies whether or not a given image is a target object (**Figure 5A**). In the first case (Train 1), the SVM is trained using an object image with various viewpoints to train the SVM, while it is trained using the object image only with a center-fixed viewpoint in the second condition (Train 2). After training, object images with various viewpoints were used for the test session (**Figure 5B, Left**). We performed this process using both invariant units and specific units.

**Figure 5.**
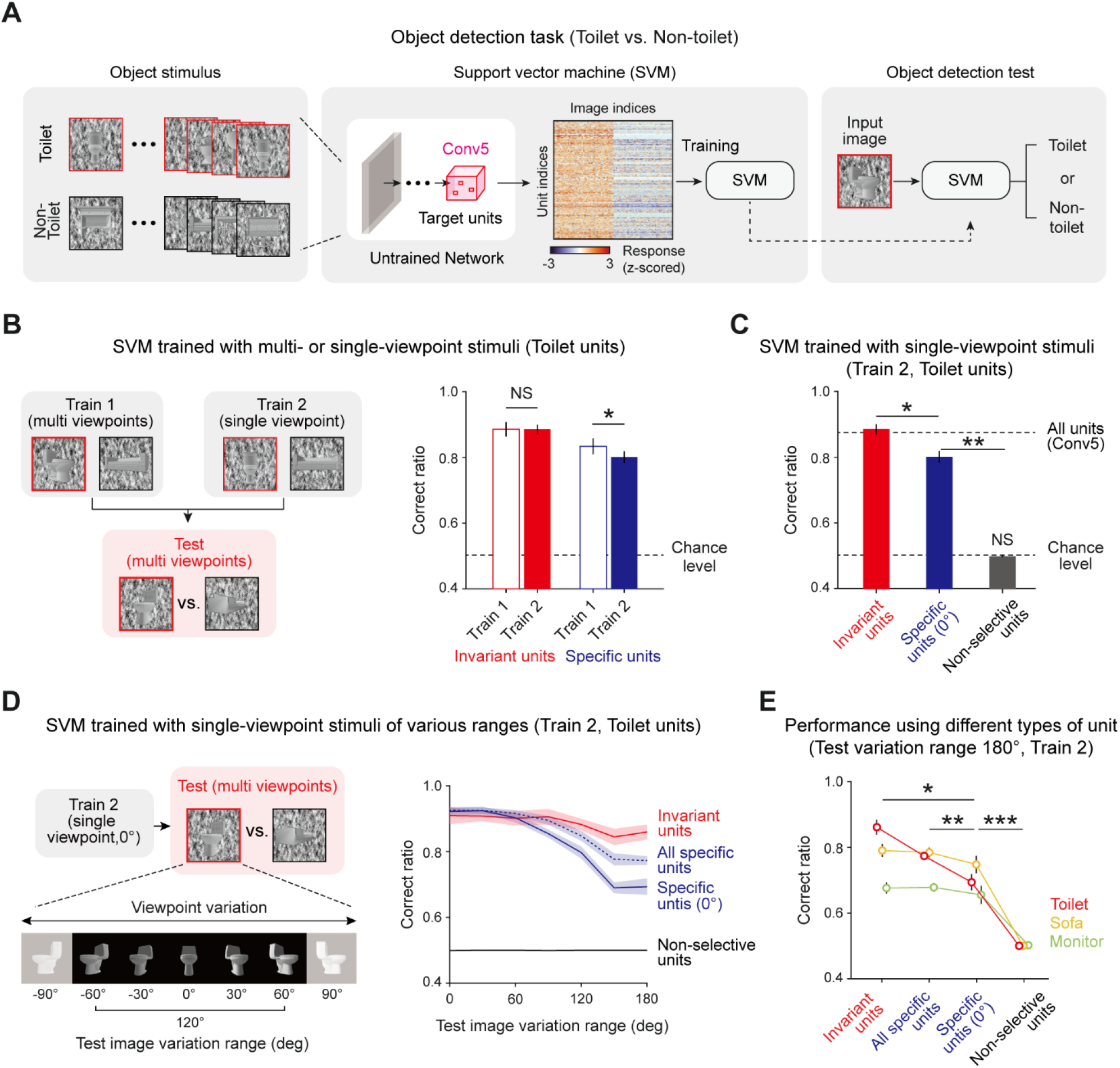
Invariantly tuned unit responses enable invariant object detection: **(A)** Overall process of the object-detection task and SVM classifier using the responses of object-selective units. To train the SVM classifier, 30 images of a target object and 30 images of a non-target object were used. Among the 60 images, 40 images were randomly sampled for training and the remaining 20 were used to test the task performance. The responses of untrained network units for these images are obtained and used to train and test the SVM to classify whether or not the given image is the target object. **(B)** Train 1 uses object images with various viewpoints to train the SVM, while Train 2 uses object images only with a center-fixed viewpoint. For the test SVM, object images with various viewpoints are used. **(C)** Performance of the Train 2 method using invariant units, specific units, and non-selective units. The upper dashed line represents the performance when all units in Conv5 are used, and the lower dashed line indicates the chance level of the task. **(D)** The task in various test-image-variation-range conditions. The test images were randomly sampled within the given viewpoint variation range. The Train 2 performances of invariant units, all specific units, specific units only with 0° preferred, and non-selective units were assessed. **(E)** Comparison of performances across the types of object units. The performance for each case was measured using test images with a 180° variation range. Shaded areas and error bars indicate the standard deviation of 20 random networks.

We found that the performances of the SVM using invariant units only and those of the SVM using specific units only are noticeably different (**Figure 5B, Right, Invariant**, n = 20, two-sided rank-sum test, NS, P = 0.735; **Specific**, n = 20, two-sided rank-sum test, *P < 0.001). Invariant units show the same level of performance regardless of training with various viewpoints, implying that the information given by invariant units is sufficient to detect images with varying viewpoints. In contrast, specific units show significantly lower performance outcomes when trained only with a fixed viewpoint (Train 2). Hence, we investigated in more depth the performances of SVMs trained in the second condition (Train 2) using different groups of units (**Figure 5C**, n = 20, two-sided rank-sum test, invariant vs. specific, *P < 10^−7^; specific vs. chance level, **P < 10^−7^; non-selective vs. chance level, NS, P = 0.116). First, the SVM using the responses of invariant units shows significantly high performance compared to the SVM using the responses of specific units. Second, the SVM using invariant units shows the same level of performance compared to when it is trained with all units in the Conv5 layer, implying that the invariance of the untrained network mainly relies on invariant units. To confirm that invariance enables invariant object detection in a wide variation range, we tested this concept in different viewpoint-variation ranges (**Figure 5D, left**). When the viewpoint variation range in the test set becomes wider, the SVM using specific units rapidly loses its performance capabilities, whereas the SVM using invariant units maintains its high-performance outcomes (**Figure 5D, right**). Interestingly, the SVM using all specific units with different preferred angles outperforms the SVM using specific units only with the same preferred angle. This trend was also observed using other object-selective units (**Figure 5E**, n = 20, two-sided rank-sum test, invariant vs. all specific, *P < 0.05; all specific vs. specific with center view, **P < 0.01; specific with center view vs. non-selective, ***P < 10^−6^). These results demonstrate that invariance in an untrained network enables an object-detection task with images with various viewpoints for a wide variation range, even without data-augmented learning.

## 3 Discussion

We showed that selectivity to various object emerges in randomly initialized networks and that this selectivity is robustly preserved even as the viewpoint changes significantly in the complete absence of learning. Furthermore, we found that the invariant tuning property can arise solely from the distribution of weights in feedforward projections. These results suggest that the statistical complexity of hierarchical neural network circuits allows the initial development of selectivity as well as invariance to various objects across a wide range of transformations.

Our results imply that innate invariance of object selectivity can arise from random feedforward projection, but this does not mean that there is no effect of experience on the development of this function. In fact, observations in various animals support the contention that this invariant function is affected by visual experience. In pigeons, the ability to detect objects across different variations in the viewing conditions is enhanced gradually during the visual training process (Watanabe, 1999). In the monkey, size-invariant object representation is reshaped by unsupervised visual experience (Li and DiCarlo, 2010). Considering the above neurological and behavioral evidence, at an early developmental stage, the innate invariance of object selectivity arises from the structure of neural circuits, and this function can be refined by visual experience during the subsequent developmental process. Specifically, repeated experience with a particular object under various viewpoints will further strengthen the existing selectivity and synaptic weights according to a biologically observed learning rule, such as Hebbian learning. In our model, the invariance of object selectivity can also be more robust and have a widened effective range by visual experience.

Although the current study investigated only viewpoint invariance, we anticipate that invariance to other types of image transformations, such as position, size, and rotation, can also emerge spontaneously in untrained neural networks. Previous studies using an untrained DNN provide supporting evidence. Kim *et al*. (Kim et al., 2021) showed that number selectivity spontaneously emerge in a randomly initialized DNN. Here, number selective units, defined as units that selectivity respond to only numbers of dots in images and respond invariantly to other visual features (e.g., locations, sizes, and convex hulls of dots), contain invariant characteristics of neural tuning. This implies that invariant tuning to various image transformations can arise in untrained neural networks. In addition, Baek *et al*. (Baek et al., 2021) found that face-selectivity can emerge initially and that this tuning shows invariant representation to position, size, rotation, and viewpoint variations to face images. Based on the above results, we expect that various types of invariance of selectivity to objects as well as faces can spontaneously arise in completely untrained neural networks.

We proposed a method of generating invariance without learning, in contrast to previous approaches that implement the same function by relying on a massive training process. In the machine learning field, invariant object recognition has been implemented by learning a great many images. To learn invariant object features, the data-augmentation method is often applied (Simard et al., 2003; O’Gara and McGuinness, 2019; Shorten and Khoshgoftaar, 2019), which generates images with variations through linear transformation, such as positional shifts, rotation, and flipping. However, data augmentation is inefficient in terms of the computational cost. One study that examined changes of the accuracy and training time by data augmentation (O’Gara and McGuinness, 2019) found that twice the learning time is required to slightly improve the accuracy by introducing data augmentation. Thus, our findings can provide clues for addressing the limitations of the data augmentation method to implement invariant functions. By finding selective units in initially randomized networks and applying a training algorithm (Zhuang et al., 2021) toward strengthening innate selectivity and invariance, we expect to reduce the computational cost of implementing invariant object recognition.

In summary, we conclude that invariance of object selectivity can arise from the statistical variance of randomly wired bottom-up projections in untrained hierarchical neural networks. Our findings may provide new insight into the developmental mechanism of innate cognitive functions in biological and artificial neural networks.

## 4 Materials and Methods

### 4.1 Untrained AlexNet

Currently, deep neural network (DNN) models, which have a biologically inspired hierarchical structure, provide an effective approach for investigating functions in the brain (Paik and Ringach, 2011; DiCarlo et al., 2012; Yamins and DiCarlo, 2016; Sailamul et al., 2017; Baek et al., 2020, 2021; Jang et al., 2020; Kim et al., 2020, 2021; Park et al., 2021; Song et al., 2021). Several studies have reported that DNNs trained to natural images can predict the neural responses of the monkey inferior temporal cortex (IT) (Cadieu et al., 2014; Yamins et al., 2014), known as the area for object recognition. Furthermore, a previous study by the authors found that face-selectivity can arise without experience using a randomly initialized DNN (Baek et al., 2021).

Following earlier work, we used a randomly initialized (untrained) AlexNet (Krizhevsky et al., 2012) consisting of feature extraction and classification layers. AlexNet extracts the features of the input image from five convolutional layers and a pooling layer. It uses a rectified linear unit (ReLU) as an activation function. This activation function allows us to investigate nonlinear activity of the type that similarly occurs in the human brain. To randomly initialize the AlexNet, we used standard randomizing method (LeCun et al., 1998). For each filter, each weight was randomly drawn from a Gaussian distribution with a zero mean and the standard deviation set to the square root of the unit number of the previous layer. With this method, we can generate an untrained state of a neural network that balances the strength of the input signal across the layers.

### 4.2 Viewpoint-controllable object stimulus renderer

There are a few well-known objects image datasets, such as ImageNet (Russakovsky et al., 2015), which are often used in DNN studies. However, this image dataset is not sufficient for investigating the effects of viewpoint variance quantitatively. Also, generally used image datasets do not control for low-level features such as luminance, contrast, position, and intra-class image similarity. For this reason, we developed a viewpoint-controllable and low-level feature-controlled object stimulus renderer.

ModelNet10 (Wu et al., 2015), a publicly available 3D object dataset which contains ten different object classes with aligned orientations, was used to render the stimulus in our study (we used only nine object classes due to an insufficient number of CAD files). Each CAD file is converted to an image at a given horizontal viewpoint using the object render. After capturing the object, the renderer generates a phase-scrambled background image. Using a sample natural image, it scrambles the phase of the given natural image in the Fourier domain and returns it to the original space. These phase-scrambled backgrounds are often used in human fMRI studies to exclude the effects of the background in visual processing (Stigliani et al., 2015). For the object images and phase-scrambled backgrounds, the overall pixel intensity is normalized in each case to have an identical intensity distribution (Pixel_mean_ = 127.5, Pixel_std_ =51). Using this renderer, we generated various viewpoint object stimulus sets in which low-level features are properly calibrated.

### 4.3 Stimulus dataset

We prepared four types of visual stimulus datasets specialized to each task. (1) Object dataset (**Supplementary Figure 1A)**: This set was used to find units that selectively respond to a particular object class. It contains nine object classes (bed, chair, desk, dresser, nightstand, monitor, sofa, table, and toilet), and 200 images are prepared in each object class. To render the images of the object dataset, the viewpoint variation angle was randomly set between −30° and +30°. In the object dataset, brightness and contrast of the images are precisely controlled to be equal across object classes **(Supplementary Figure 1B and C)**. In addition, the intra-class similarity of the images in each object category was calibrated at a statistically comparable level **(Supplementary Figure 1D)**. (2) Viewpoint dataset **(Supplementary Figure 3A)**: This set was used to test the viewpoint-invariant characteristics of the object-selective units. This dataset consists of 13 subsets which have different viewpoints from −180° to +180° on a linear scale. It contains 250 different object identities in an object class. Among them, 200 object identities are identical to those used in the object dataset. They were used to analyze the viewpoint-invariant characteristics of the object-selective units quantitatively. The remaining 50 object identities were used not to find object-selective units but to distinguish object-selective units with or without viewpoint invariance. In the viewpoint dataset, the luminance and contrast are also controlled **(Supplementary Figures 3B and C)**. (3)SVM dataset: This set was used to train and test the SVM that performs the object-detection task. It contains 60 different object identities in an object class, which were not used for finding object-selective units. Specifically, it consists of 18 subsets with different viewpoint variations ranging from 0° to 180°. For example, a subset with a 180° viewpoint variation range contains images that show different viewpoints of objects within −90° and +90°.

### 4.4 Analysis of responses of the network units

Using the totally untrained AlexNet, we measured the responses of the target layer for each designed stimulus. For each response from the target convolution layers, each unit of an activation map was separately recorded for different classes of the stimulus. Based on our previous study, object-selective units were defined as units that showed a significantly greater mean response to target object images compared to those of non-target object images (P < 0.001, two-sided rank-sum test). To analyze the responses of each unit, it was necessary to regularize the raw response. To normalize the raw response, we used the z-scoring method. Furthermore, we used a trick in the z-score in that we subtracted 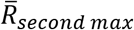 from 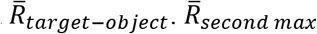 indicates the response for an object class that leads to the second maximum response for that unit. Therefore, if the z-scored response is higher than zero, our unit shows a higher raw response to the target object than to the second maximum object, indicating selectivity.

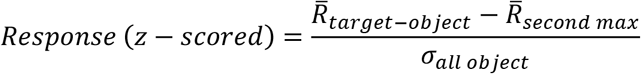

To quantify the degree of tuning, an object selectivity index (OSI) of a single unit was defined using the follow formula. This index is modified from the face-selective index (FSI), which defined in previous experimental research (Aparicio et al., 2016).

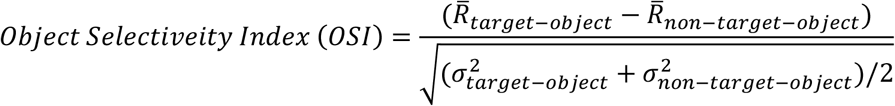

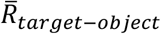 is the average response to target-object images and 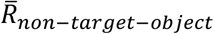 is the average response to all non-target-object images. A higher OSI indicates fine tuning and an OSI of zero indicates equal responses to target and non-target object images.

Among the object-selective units, we defined a viewpoint-invariant unit as a unit for which the response was not significantly different (one-way ANOVA, P > 0.05) for all viewpoint classes. Similarly, viewpoint-specific units are defined as a unit for which the response was significantly high for one preferred viewpoint class (one-way ANOVA, P < 0.05).

To measure the invariant index quantitatively, we calculated the inverse of the standard deviation of the average responses for images within each viewpoint class.

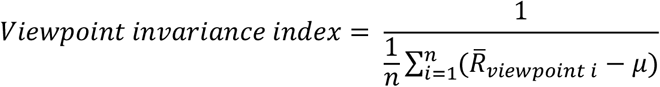

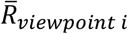 is the average response to a viewpoint class and *μ* is average response for all viewpoint classes. is total number of viewpoint classes.

### 4.5 Preferred feature image (PFI) analysis

To visualize the preferred input features of target units, the receptive field was estimated by the reverse correlation method (Bonin et al., 2011) with multiple iterations. The initial stimulus set was generated using 2500 random local 2D Gaussian filters. Such stimuli were weighted by the corresponding responses and were added as an initial preferred feature image. Then, to detect the preferred feature more accurately, we calculated the PFI iteratively; the PFI of the next iteration was calculated by a new stimulus set consisting of the summation of the current PFI and the 2500 random local 2D Gaussian filters. We repeated 100 iterations and obtained the final PFI.

### 4.6 Connectivity analysis

To investigate the connectivity between object-selective units across convolutional layers, we backtracked projections of the units from the source layer (Conv4) to the projection layer (Conv5). This backtracking process is opposite of the group convolution process. To backtrack the origin of a unit in the projection layer, we investigated all connected weights and units in the source layers.

To measure the degree of homogeneity in the input projection weight to a single target unit, the homogeneous index was defined as

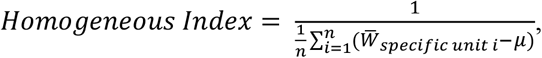

where 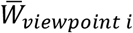 is the average weight from specific units in the source layer to a unit in the projection layer and *μμ* is the average weight from all specific units. *RR* is the total number of viewpoint-specific units with different preferred angles. To compare the unbiased properties of specific and invariant units, we normalized the homogenous index so that the average index value of viewpoint-specific units reaches unity.

### 4.7 Object-detection task

To validate viewpoint-invariant object-selectivity that spontaneously emerges in an untrained DNN, we trained a support vector machine (SVM) using the responses of object-selective units with two types of training. For Train 1, target-object (n = 40) or non-target-object (n = 40) images, which shows different viewpoints of objects within a range of −60° and +60° were randomly presented to the networks, and the observed responses of the Conv5 layer were used to train the SVM. For Train 2, most of the processes are nearly identical compared Train 1, but the only difference is in how the train images are presented. We prepared target-object and non-target object images without viewpoint variation (front-view only). After training the SVM, we investigated the performance with the responses of object-selective units for a stimulus with viewpoint variation. Here, target-object (n = 20) or non-target-object (n = 20) images were also randomly presented to the networks, and the responses from the Conv5 layer was used to test the SVM.

## 5 Data Availability Statement

The stimulus datasets and the MATLAB codes for this study are available at https://github.com/vsnnlab/Invariance.

## 6 Author Contributions

S.-B.P. conceived of the project. J.C., S.B. and S.-B.P. designed the model. J.C. performed the simulations. J.C. and S.B. analyzed the data. J.C., S.B. and S.-B.P. wrote the manuscript.

## 11 Supplementary Figures

**Supplementary Figure 1.**
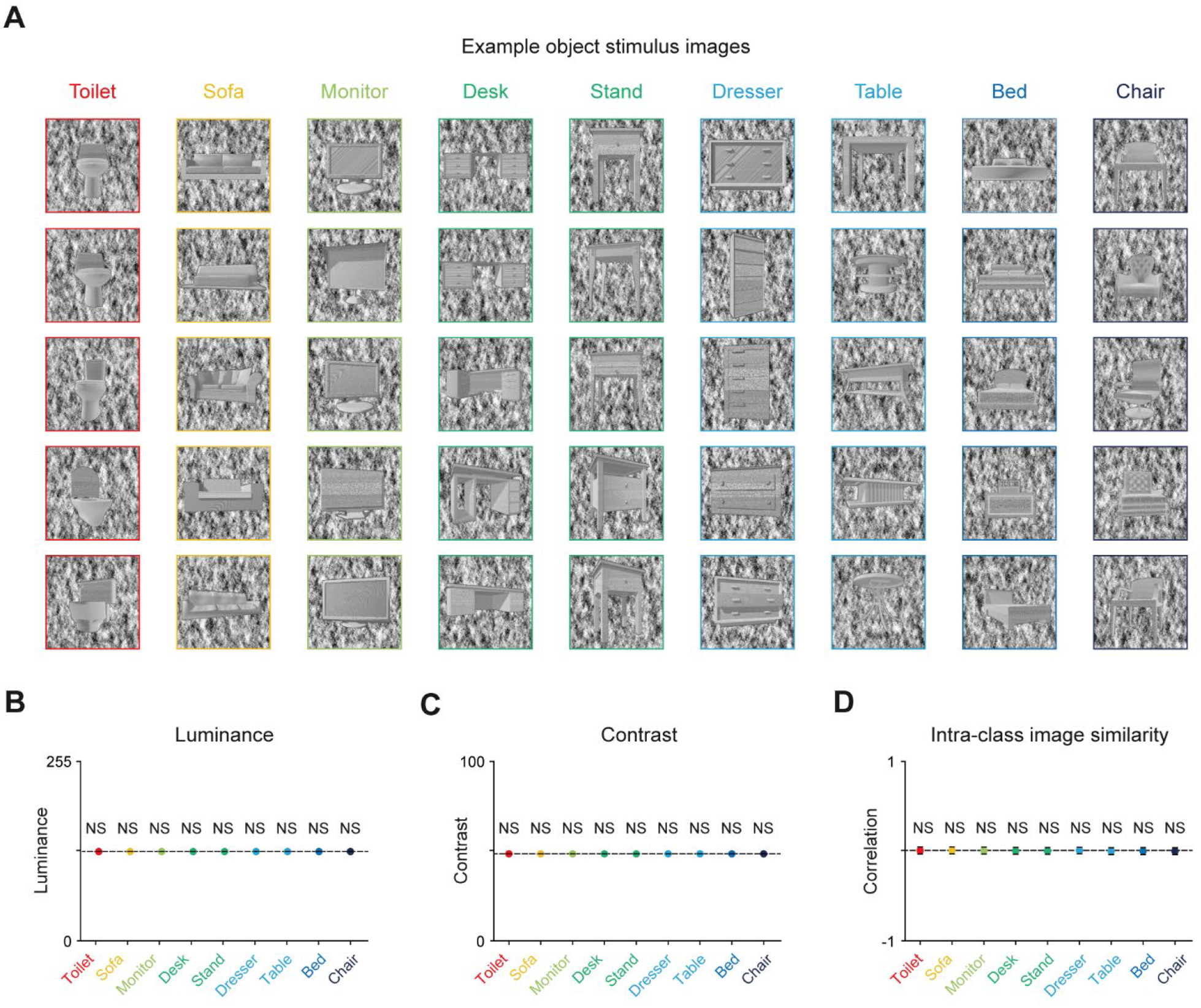
Low-level feature-controlled object image stimulus: **(A)** Example images of the object dataset. An object stimulus renderer was used to create object images with different viewpoints. Object stimulus images were selected and adapted from a publicly available dataset that contains nine object classes. Each object set contains 200 different object identities rendered with viewpoint variations of ±30°. The original CAD models are available at https://modelnet.cs.princeton.edu/. **(B)** Controlled luminance of stimulus images (n = 200, one-sided sign rank test, NS, P > 0.829). The luminance of an image was measured as the mean of the pixel intensity. **(C)** Controlled contrast of object stimulus. The contrast of an image was measured as the standard deviation of the pixel intensity. The luminance and contrast for every stimulus are set to be identical. (n = 200, one-sided sign-rank test, NS, P = 0.507). **(D)** Controlled intra-class similarity of the object stimulus. The intra-class similarity was measured as the image correlation between the images in each class (n = 19,900, two-sided rank-sum test, NS, P = 0.767). Error bars indicate the standard deviation (n = 19,900).

**Supplementary Figure 2.**
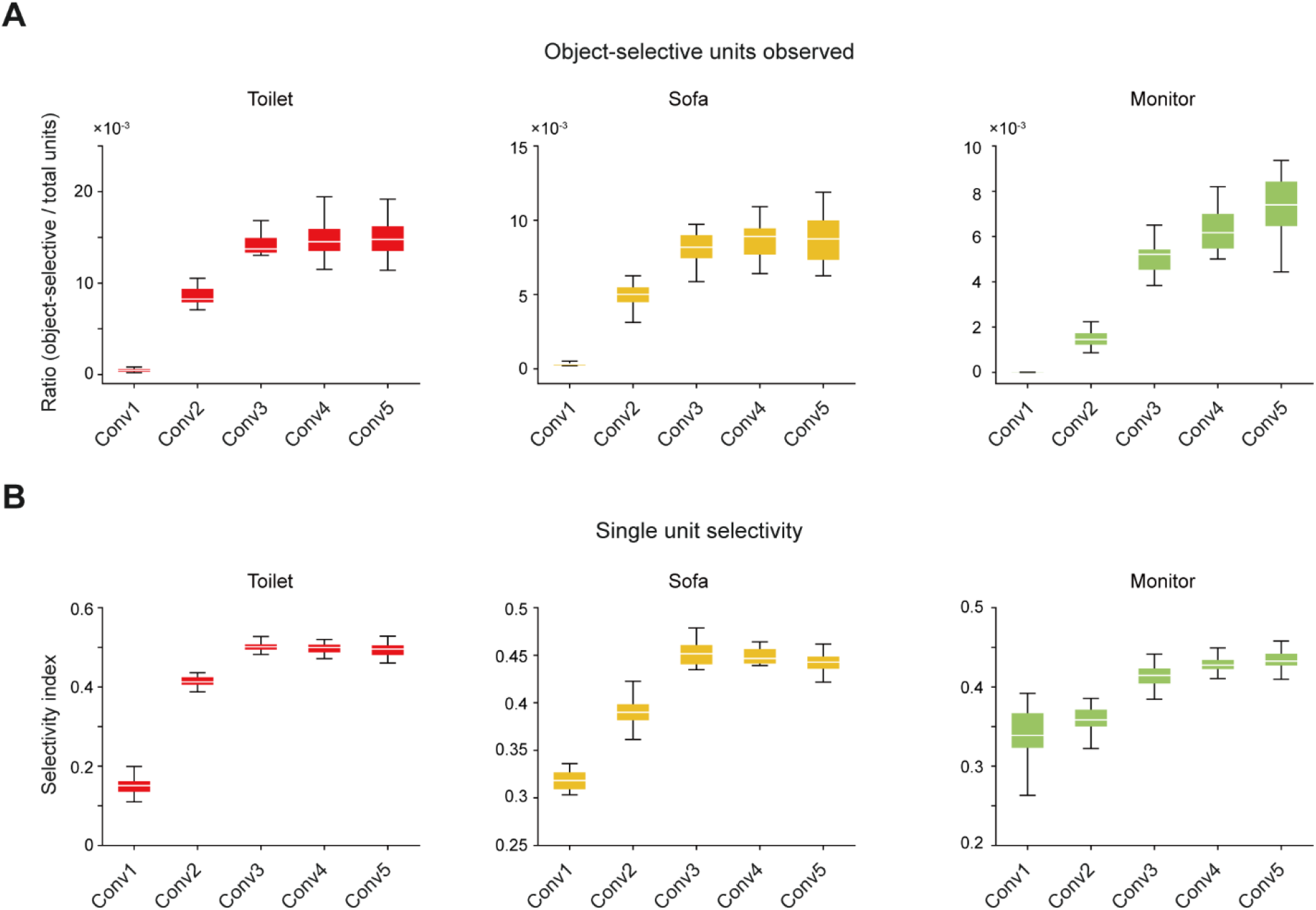
Layer-dependent object selectivity across the hierarchal layer in untrained networks: **(A)** Number of selective units across convolutional layers in untrained networks (n = 20). **(B)** Object-selective index of single units (n = 20) across the convolutional layers. Box plots indicate the inter-quartile range (IQR between Q1 and Q3) of the dataset. White lines depict the median and whisker plots indicate the rest of the distribution (Q1 − 1.5*IQR, Q3 + 1.5*IQR).

**Supplementary Figure 3.**
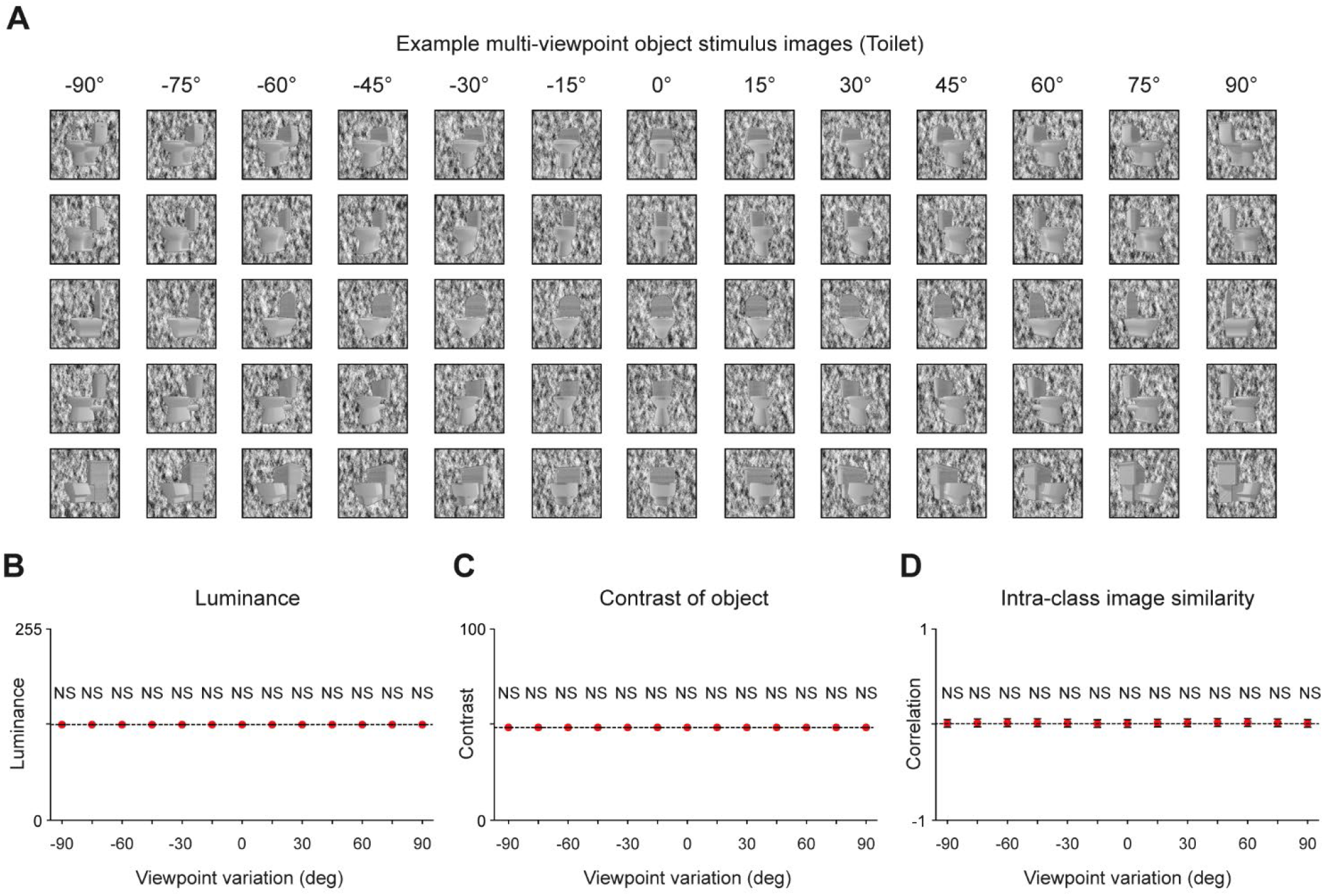
Low-level feature-controlled object image stimulus with various viewpoints: **(A)**Example images of the viewpoint dataset. Using the viewpoint-controllable object stimulus renderer, a viewpoint dataset that contains 13 viewpoint classes from −180° to +180° was generated. **(B)**Controlled luminance of stimulus images (n = 200, two-sided rank-sum test, NS, P > 0.816). **(C)** Controlled contrast of stimulus images. The luminance and contrast for every stimulus are set to be identical (n = 200, two-sided rank-sum test, NS, P = 0.485). **(D)** Controlled intra-class similarity of the object stimulus. The intra-class similarity was measured as the image correlation between the images in each class (n = 19,900, two-sided rank-sum test, NS, P = 0.492). Error bars indicate the standard deviation (n = 19,900).

**Supplementary Figure 4.**
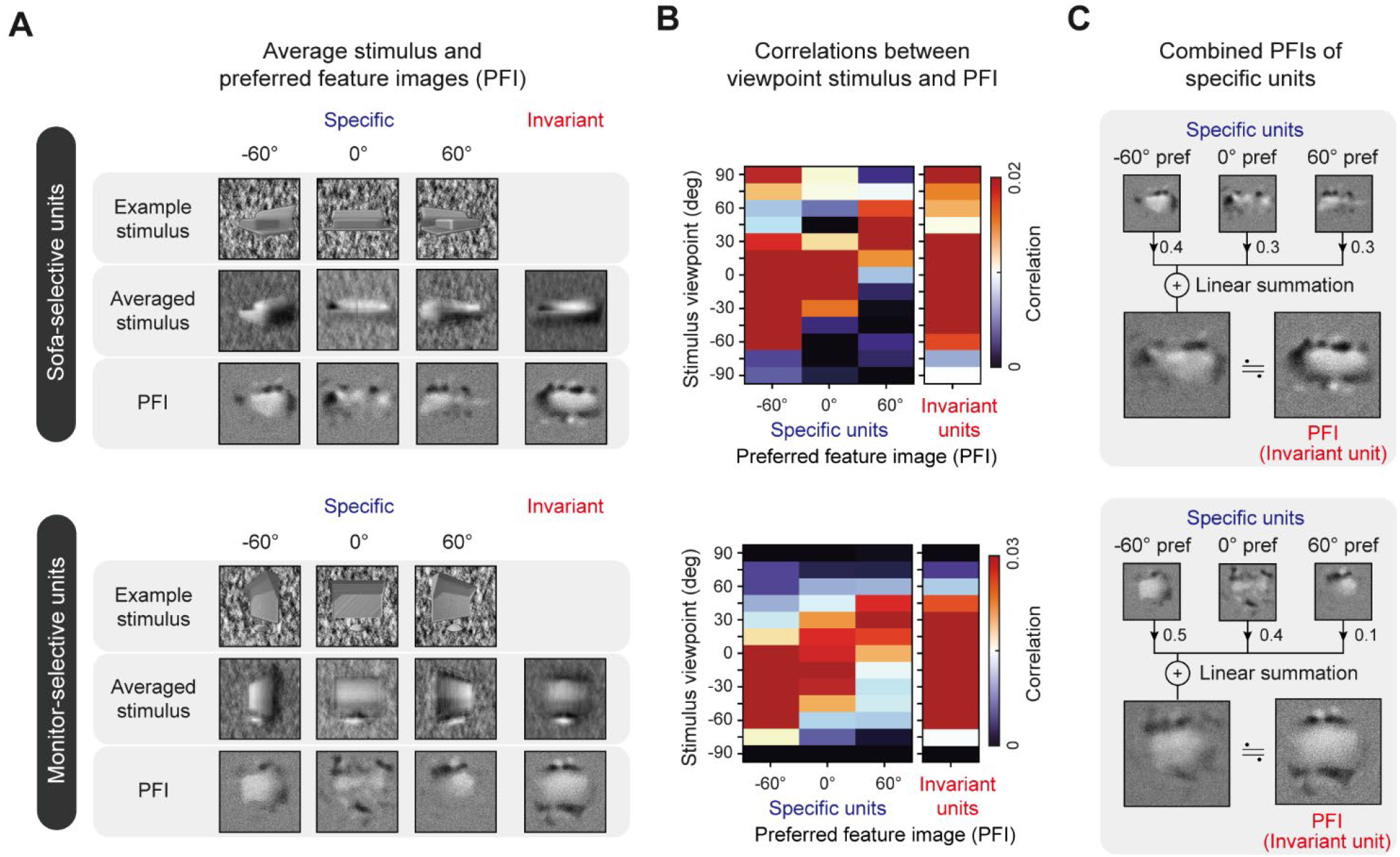
Preferred feature image (PFI) analysis for specific and invariant units: **(A)** Example PFIs of invariant and specific units. An example stimulus and the average stimulus are also presented together for a visual comparison. **(B)** Correlation between the PFIs of each type of unit and stimulus with different angles. **(C)** Linear combination of PFIs of specific units showing the highest correlation with the PFIs of invariant units.

**Supplementary Figure 5.**
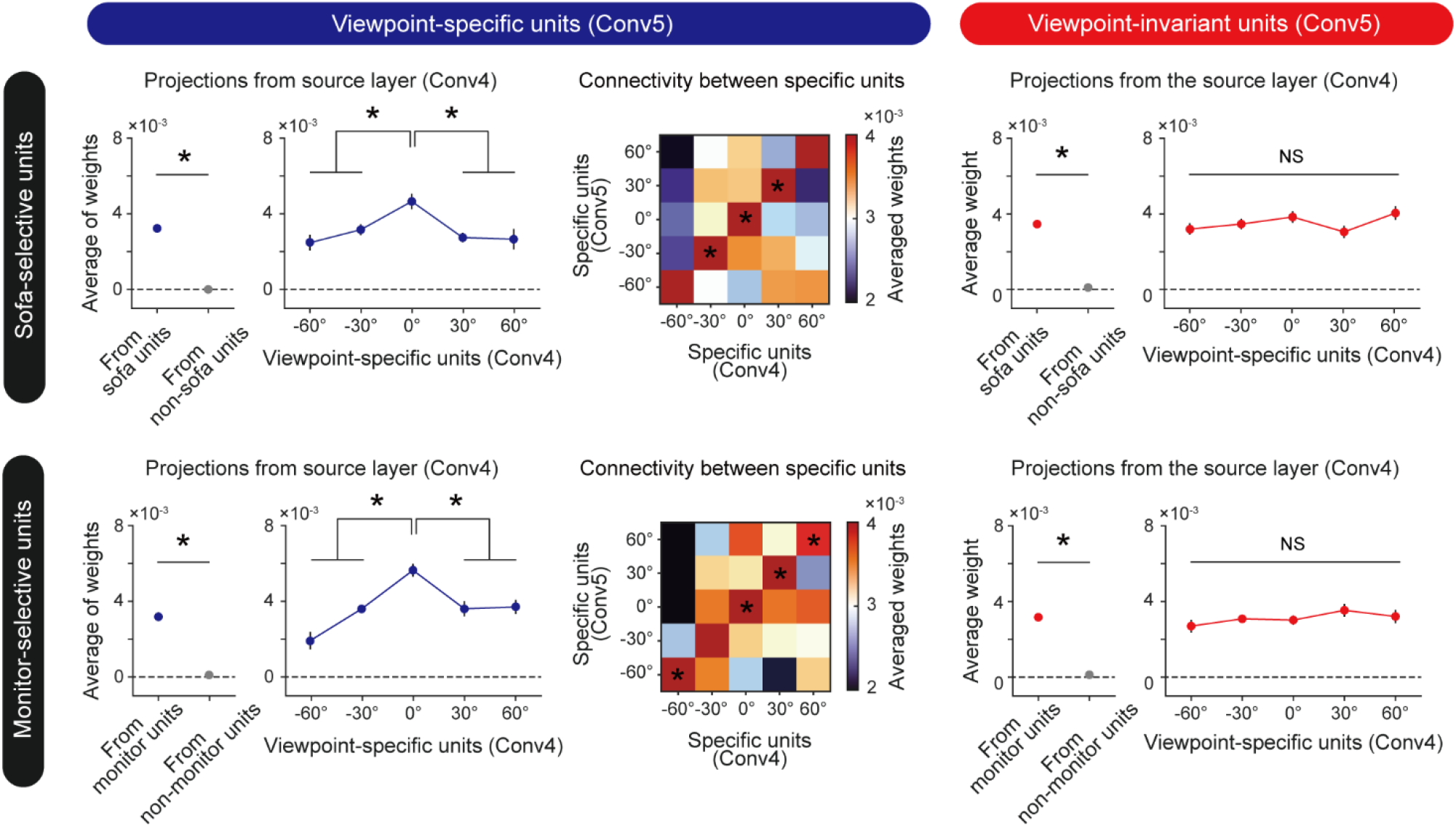
Emergence of viewpoint invariance: In each panel, the leftmost graph demonstrates how strongly a target unit is connected to each type of unit in the source layer (n = 20, two-sided rank-sum test; specific sofa unit, *P < 10^−7^; invariant sofa unit, *P < 10^−7^; specific monitor unit, *P < 10^−7^; invariant monitor unit, *P < 10^−7^). The graph next to the leftmost graph indicates how strongly a unit in the target layer is connected to specific units with different angles in the source layer (n = 20, one-way ANOVA; specific sofa unit, *P < 10^−3^; invariant sofa unit, NS, P = 0.132; specific monitor unit, *P < 0.05; invariant monitor unit, NS, P = 0.991). Error bars indicate the standard error of 20 random networks. The heatmap, located at the center between two panels, shows how strongly the specific units with various preferred angles are connected between the source and target layers (n = 20, one-way ANOVA; specific sofa unit, *P < 0.001; specific monitor unit, *P < 0.05).

**Supplementary Figure 6.**
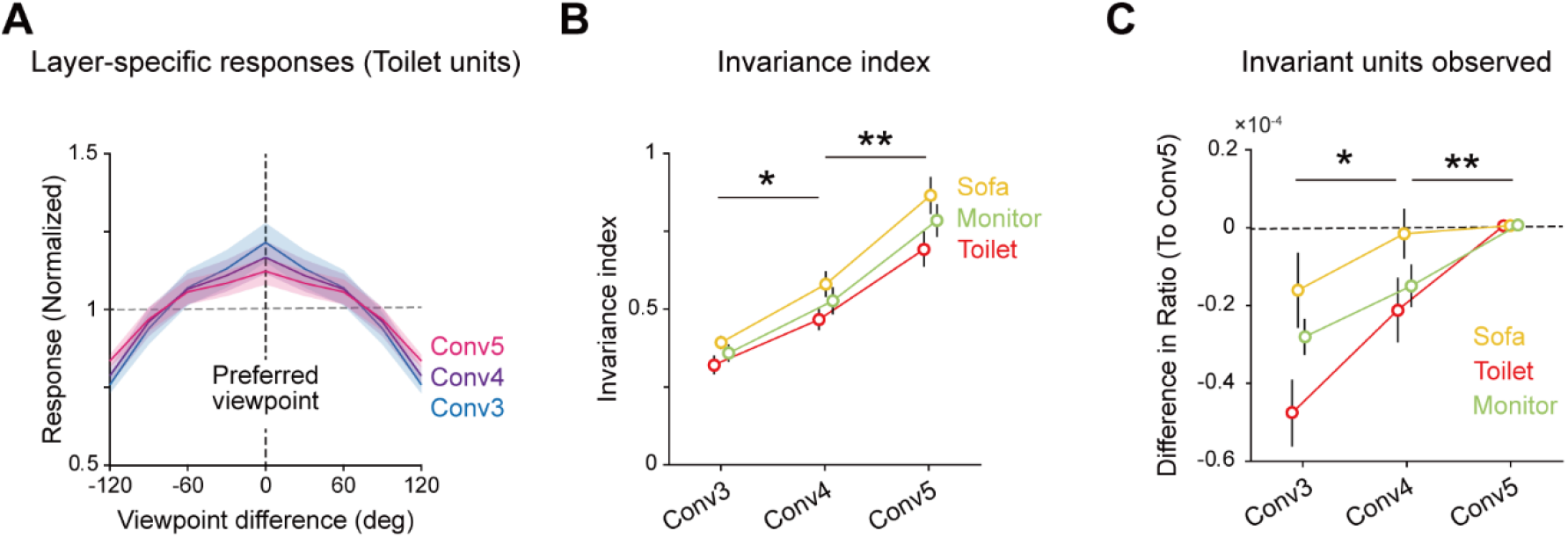
Layer-dependent invariance across the hierarchal layer in untrained networks: **(A)** Normalized average tuning curve of toilet-selective units in each convolutional layer. Note that the tuning curves become flatter as the layer deepens, demonstrating increased invariance of tuning. The shaded area indicates the standard error of all units in each layer. **(B)** Invariance index of object units across convolutional layers. **(C)** The ratio of invariant units across convolutional layers. Circles and error bars correspondingly indicate the mean and the standard error of 20 random networks in **(B)** and **(C)**.

